# State-dependent cortical unit activity reflects dynamic brain state transitions in anesthesia

**DOI:** 10.1101/2020.06.13.150235

**Authors:** Heonsoo Lee, Shiyong Wang, Anthony G. Hudetz

## Abstract

How anesthesia affects cortical neuronal spiking and information transfer could help understand the neuronal basis of conscious state. Recent investigations suggest that global state of the anesthetized brain is not stationary but changes spontaneously at a fixed level of anesthetic concentration. How cortical unit activity changes with dynamically transitioning brain states under anesthesia is unclear. We hypothesized that distinct cortical states are characterized by distinct neuronal spike patterns. Extracellular unit activity was measured with sixty-four-channel silicon microelectrode arrays in cortical layers 5/6 of primary visual cortex of chronically instrumented, freely moving male rats (N = 7) during stepwise reduction of the anesthetic desflurane (6, 4, 2, and 0%). Unsupervised machine learning applied to multi-unit spike patterns revealed five distinct brain states of which four occurred at various anesthetic concentrations and shifted spontaneously. In deeper anesthesia states, the number of active units and overall spike rate decreased while the remaining active units showed increased bursting (excitatory neurons), spike timing variability, unit-to-population correlation and unit-to-unit transfer entropy, especially among putative excitatory units, despite the overall decrease in transfer entropy. A novel desynchronized brain state with increased spike timing variability, entropy and electromyographic activity that occurred mostly in deep anesthesia was discovered. These results provide evidence for distinct unit activity patterns associated with spontaneous changes in local cortical brain states at stationary anesthetic conditions. The appearance of a paradoxical, desynchronized brain state in deep anesthesia contends the prevailing view of monotonic dose-dependent anesthetic effects on the brain.

**SIGNIFICANCE STATEMENT:** Recent studies suggest that spontaneous changes in brain state occur under anesthesia. However, the spiking behavior of cortical neurons associated with such state changes has not been investigated. We found that local brain states defined by multi-unit activity had non-unitary relationship with the current anesthetic level. A paradoxical brain state displaying asynchronous firing pattern and high electromyographic activity was found unexpectedly at high-dose anesthesia. In contrast, the synchronous fragmentation of neuronal spiking appeared to be a robust signature of the state of anesthesia. The findings challenge the assumption of monotonic, anesthetic dose-dependent behavior of cortical neuron populations. They enhance the interpretation of neuroscientific data obtained under anesthesia and understanding of the neuronal basis of anesthetic-induced state of unconsciousness.

## INTRODUCTION

Recent studies of large-scale brain activity found that multiple brain states appear at a constant anesthetic concentration and conversely, one brain state can be observed at different anesthetic levels (Chander et al., 2014; Hudson et al., 2014; Hudson, 2017; Li et al., 2019). The degeneracy in the relationship between brain state and anesthetic concentration suggests that the neuronal network spontaneously shifts between two or more transient attractors (metastability) or switches multiple stable attractors via external perturbation or noise (multistability) (Hudson et al., 2014; Hudson, 2017; Li et al., 2019).

Despite these observations, most studies of unit activity assume a one-to-one relationship between brain state and anesthetic concentration and investigate dose-dependent changes of neuronal activity (Vizuete et al., 2012; Sellers et al., 2013; Vizuete et al., 2014). This would be surprising and suggests that the spiking dynamics of individual unit activities in different brain states under anesthesia has been poorly explored. Detailed information about the spiking dynamics during shifting brain states is arguably important for interpreting neuroscientific data obtained under anesthetized conditions and to understand the neuronal mechanisms of changing states of consciousness.

In an attempt to fill this gap of knowledge, we measured single unit spiking patterns of neuronal populations in chronically instrumented rodents subjected to multiple levels of anesthesia and applied machine learning to identify brain states independent of the actual anesthetic concentration. We hypothesized that brain states identified by specific features of population (multi-unit) activity will show degeneracy in the relationship with anesthetic concentration and that these states will be characterized by distinct spike activity patterns.

## METHODS

### Electrode implantation

The study was approved by the Institutional Animal Care and Use Committee in accordance with the Guide for the Care and Use of Laboratory Animals of the Governing Board of the National Research Council (National Academy Press, Washington, D.C., 2011).

Eight adult male Long-Evans rats (300-350 g weight) were housed in a reverse light-dark cycle room for 5-7 days prior to surgical implantation. Ad libitum access for food and water was provided while the animals remained in the room for the duration of the experiment. A multi electrode array consisting of 64-contact silicon probes (shank length 2 mm, width 28-60 μm, probe thickness 15 μm, shank spacing 200 μm, row separation 100 μm, contact size 413 μm^2^; custom design a8×8_edge_2mm100_200_413, Neuronexus Technologies, Ann Arbor, MI) was chronically implanted in the primary visual cortex of each rat. A pair of insulated wires (PlasticsOne, Inc., Roanoke, VA), exposed at the tips, was positioned bilaterally into the nuchal muscles to record electromyogram (EMG).

A craniotomy of rectangular shape of approximately 2 × 4 mm was prepared, the exposed dura mater was resected, and the electrode array was inserted using a micromanipulator to the final position 1.6 mm below the pial surface. The perimeter was covered with silicone gel (Kwik-Sil, World Precision Instruments, Sarasota, FL). Additional sterilized stainless-steel screws were used to secure the electrode to the cranium. The assembly was embedded with Cerebond (MyNeurolab, Saint Louis, MO). Carprofren (5 mg/kg s.c. once daily) was administered for 2 and 7 days, respectively. The animals were observed for 7–10 days for any infection or other complications.

### Experimental design

One to eight days after surgery, the animals were placed in a closed, ventilated anesthesia chamber for continuous recording of extracellular potentials in dark environment. Desflurane was administrated with a stepwise decreasing concentration, 6%, 4%, 2%, and 0%. Between every concentration levels, there was a 15 minutes of transient period which allowed to reach equilibrium concentration (Fig. 1A). Each concentration level comprised of resting state and visual stimulation sessions, during which light flashes of 1 and 10 ms durations were delivered to the retina by transcranial illumination with randomized intervals (2-4 seconds). Neural response to visual flashes is beyond the scope of the study and thus the electrophysiological recordings during the visual stimulation session were not used in this study.

**Figure 1.**
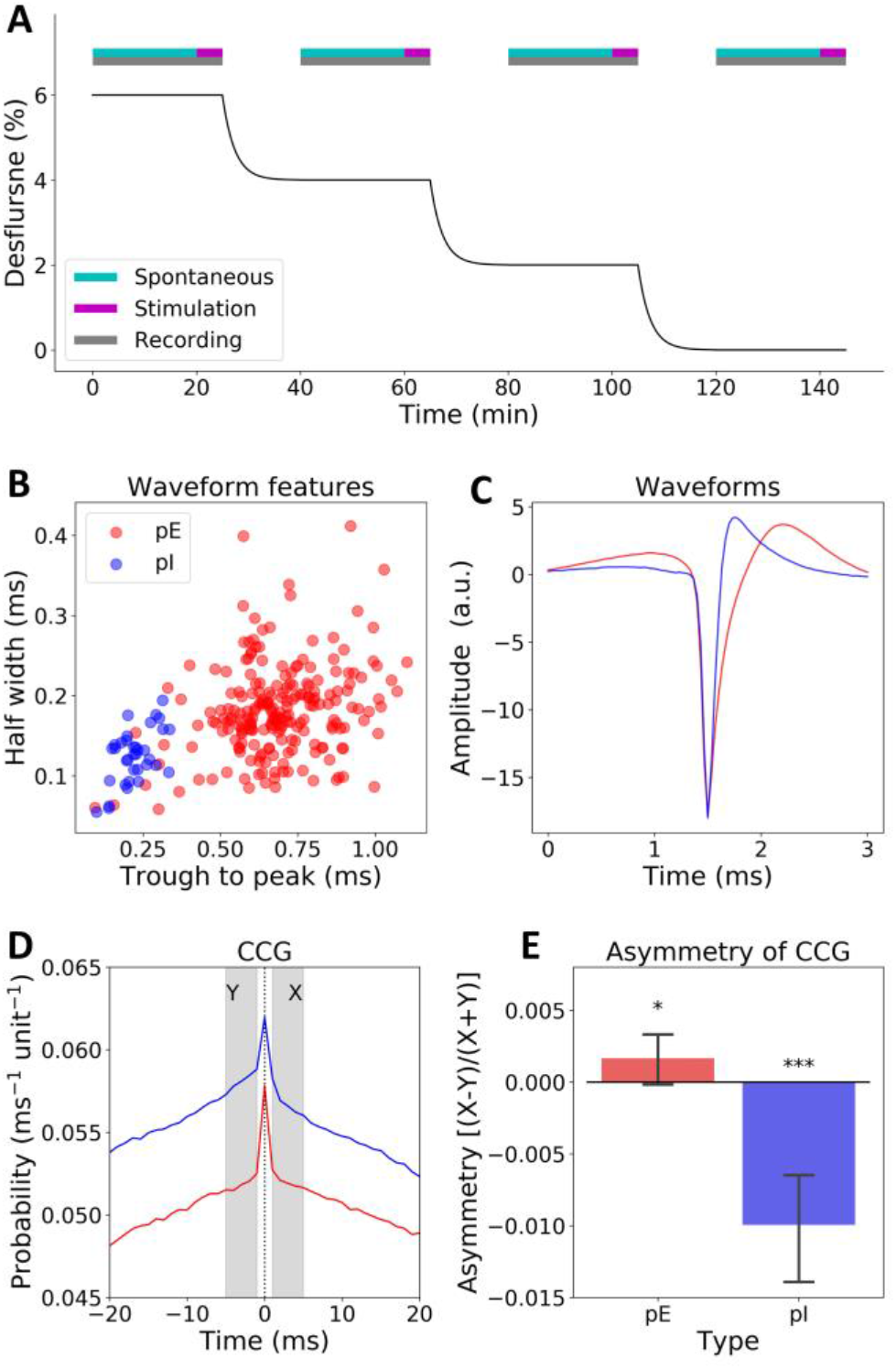
Anesthetic procedure and identification of excitatory and inhibitory units. (A) Schematic representation of the anesthesia protocol. The volatile anesthetic desflurane was administered at steady state concentrations and decreased in a stepwise manner. This study used spontaneous activity data only. (B) Putative excitatory (pE) and inhibitory (pI) units were classified based on spike waveform features and the autocorrelogram (not shown). (C) Average spike waveform of pE (n=215) and pI (n=36) units. (D) Average cross-correlogram (CCG) between single unit activity and multi-unit activity (MUA). The cross-correlogram was normalized by dividing the number of units in MUA. (E) Asymmetry of CCG confirms the identification of pE and pI units. Error bar indicates 95% confidence interval across units (*p<0.05, ***p<0.001; one-sided *t* test with Bonferroni correction).

Spontaneous activity during resting state session was recorded for twenty minutes per each desflurane level. For one experiment which was performed in the beginning of the study, only forty minutes of spontaneous activity was recorded (ten minutes per anesthetic concentration). Because all measurements of neuronal activity (spike rate, burst ratio, etc.) were quantified from 10 second epochs, ten minutes data length per desflurane concentration should not affect the final conclusions and the data was kept for the analysis. Anesthetic concentration in the holding chamber was continuously monitored (POET IQ2 monitor; Criticare Systems, Inc., Waukesha, WI). Core body temperature was maintained at 37°C by subfloor heating.

### Electrophysiological Recording and identification of single units

Extracellular potentials were recorded using SmartBox (Neuronexus Technologies, Ann Arbor, MI) at 30 kHz sampling rate. The data were used for both detecting unit activities (high frequency components; > 300 Hz) and for local field potentials (low frequency components; < 100 Hz). To investigate unit activities, the sixty-four signals were median-referenced. For every time stamp with signal amplitude larger than 10 SD, the periods ± 1 second of those time stamps were removed. The records were also visually inspected for noticeable noise episodes that were manually excluded from the analysis. One experiment was excluded from the analysis due to severe noise contamination (n = 7).

Single unit activity (SUA) was identified using the clustering software Spiking Circus, a template-based spike sorting method (Yger et al., 2018). On average, 36 ± 14 (mean ± SD) single units were obtained per animal. The SUAs were further classified into putative excitatory (pE) and inhibitory (pI) units based on the spike waveform, autocorrelogram and cross-correlogram. Units with short half-amplitude width, short trough-to-peak time, and fast-spiking pattern (a prominent peak near 10-30 ms of autocorrelogram) were manually selected as a pI (Csicsvari et al., 1998; Sirota et al., 2008) (Fig. 1B-C). The rest of the units were classified into PE. Cross-correlogram can be used to identify putative monosynaptic connections (Vizuete et al., 2012) but the chance of finding connections is small when the recording sites are relatively far from each other. As an alternative, we calculated cross-correlogram between individual units (reference unit) and multi-unit activity (MUA; the summation of SUAs), then compared the level of MUA before and after spike events of individual units (Fig. 2D). That is, our approach is based on a conjecture that pI (pE) units, on average, inhibit (excite) other units resulting in a negative (positive) asymmetry in cross-correlogram; the asymmetry of cross-correlogram was defined as (X-Y)/(X+Y), where X (Y) is the number of spike events of all the other units 1 to 5 second after (before) the spike of the reference unit. All properties of local field potential (LFP) and spikes were calculated for non-overlapping 10 second epochs by assuming stationarity over the timescale of anesthetic-induced slow oscillations and burst-suppression pattern.

**Figure 2.**
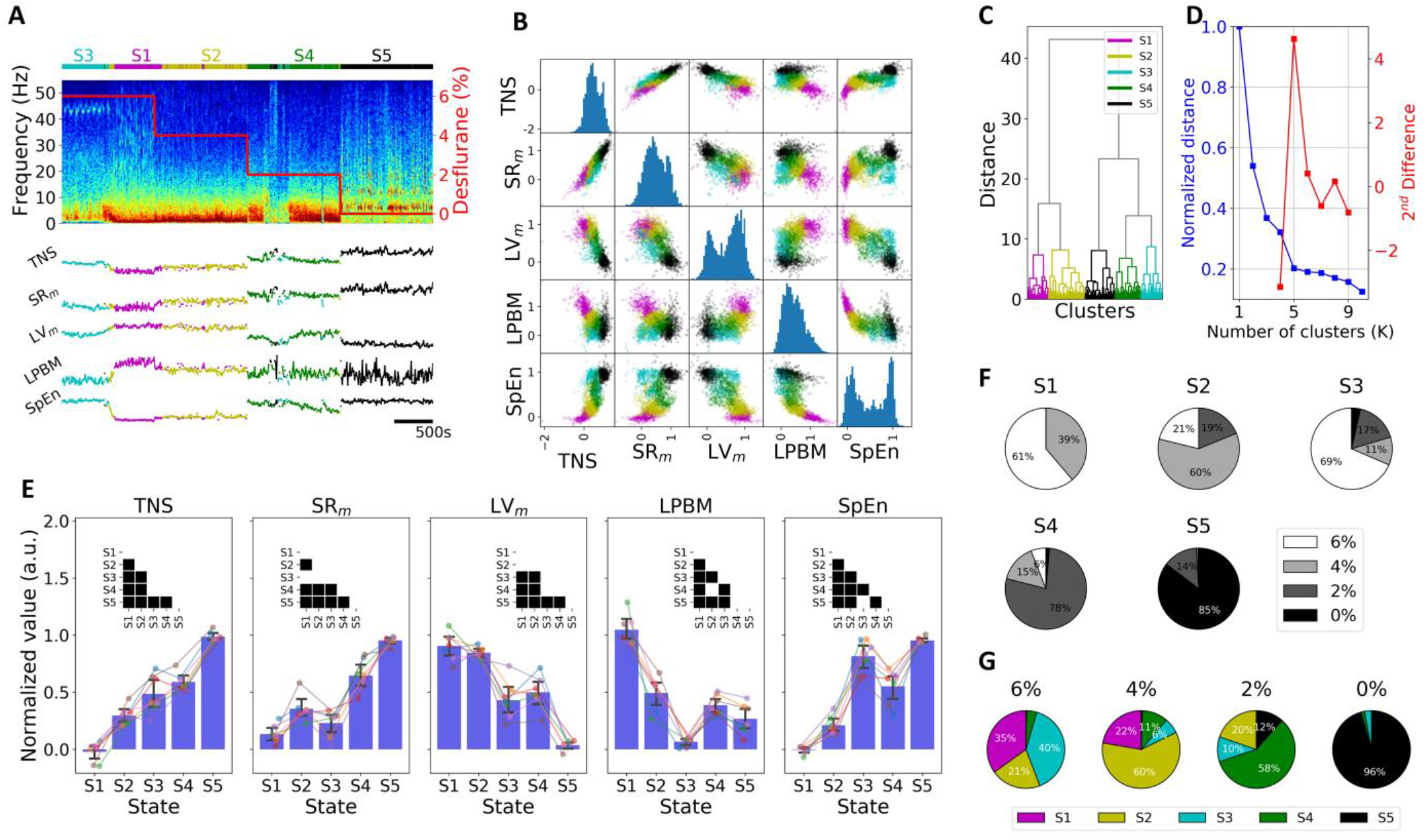
Classification of brain states. (A) Local field potential spectrogram and traces of five features from a representative animal; TNS: total number of spikes, SR_m_: mean of log-transformed spike rate, LV_m_: mean local variation, LPBM: longest period below mean, SpEn: sample entropy. Colors in the horizontal bar above the spectrogram indicate different brain states (S1 through S5) as classified by clustering. (B) Scatter and histogram plots of the five features used for state clustering from pooled data. (C) Dendrogram generated from hierarchical agglomerative clustering. (D) Elbow method suggests the optimal number of clusters as *K*=5. (E) Brain state-dependent changes of the five features. Error bar indicates 95% confidence interval across animals. The inset in each panel represents statistically significant difference between pairs of brain states. (F) Relative frequency of four anesthetic concentrations supporting each brain state; pooled data from seven animals. (G) Relative frequency of five brain states at each anesthetic concentration; pooled data from seven animals.

### Spectral analysis of LFP

LFP signals were median-referenced, and one high-quality channel was chosen for the spectral analysis. For every time stamp with signal amplitude larger than 5 SD, the periods ±1 second of those time stamps were removed. Power spectral density (PSD) of LFP in each epoch was obtained by Welch’s method; the 10 second epochs were divided into 4 second windows with 50% overlap, and time series data in each window was multiplied with Hanning window to perform the fast Fourier transform. A function “welch.py” in Python SciPy library (http://www.scipy.org) was used. The calculated PSDs from each epoch were concatenated in order to visualize the time-varying pattern of PSD (spectrogram). For the comparison of PSDs between different brain states, PSDs from epochs in each brain state were averaged.

### EMG activity

EMG signal was recorded with 1–500 Hz analog band-pass filter and 30 kHz sampling rate, and was used as a surrogate measure of the vigilance level. EMG signal was first down-sampled to 3 kHz and PSD was calculated using the same parameters with the PSD calculation of LFP signal. The PSD values with frequencies lower than 250 Hz were discarded due to cardiac artifact contamination. Next, overall EMG activity level at each of the consecutive epochs was estimated by the sum of the log-transformed PSD values. For a comparison across different animals, EMG activity was to a range between zero and one.

### Single-unit spike properties

*Spike rate (SR)* Because SR is known to follow lognormal distribution, linear-scale SR values were log-transformed (Buzsáki and Mizuseki, 2014), and averaged for nonoverlapping consecutive 10-second epochs. Zero spike rates were substituted by SR = 10^−2^ Hz before the log-transformation.

*Gini coefficient* The Gini coefficient was used to estimate the dispersion of the SR distribution. It was originally intended to represent the income or wealth disparity, and is commonly being used in measurement of inequality. For non-negative values, the Gini coefficient can theoretically range from zero to one, zero being complete equality and one being complete inequality. Gini coefficient was calculated from raw (not log-transformed) SR data, by plotting the neuronal population sorted by SR on the x-axis and cumulative SR on the y-axis (Lorenz curve, (Lorenz, 1905)). The area below the Lorenz curve of the empirical SR data (area A) is then compared to the area below the Lorenz curve of an ideal SR data (area B), in which all neurons have an equal SR value. The Gini coefficient value is finally defined as a ratio, (B-A)/B.

*Burst ratio (BR)* Two spikes with short inter-spike intervals (ISI) (<10 ms) were considered as an indication of bursting spike. BR was defined as the number of ISIs shorter than 10 ms (bursting spikes) divided by the total number of ISIs in each epoch. Units with SR < 1 Hz in each epoch were considered as inactive units and excluded from the BR calculation. BR values were log-transformed and averaged for consecutive 10-second epochs.

*Local variation (LV)* Spike timing variability, or spike irregularity was estimated by local variation (Shinomoto et al., 2003) from each spike train of SUA. LV was defined as,

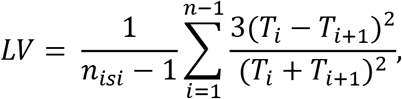

where *T*_*i*_ is the duration of *i*th ISI and *n*_*isi*_ is the number of ISIs. LV is zero for constant *T*_*i*_, and approaches one for a sufficiently long Poisson ISI sequence. LV is thought to differentiate the degree of intrinsic spiking randomness of individual neurons more effectively than the other measures, such as coefficient of variation of ISI (Shinomoto et al., 2003). LV values were not log-transformed because it did not show lognormal distribution.

*Transfer entropy (TE)* was used to estimate directional functional connectivity among individual units (Schreiber, 2000; Ito et al., 2011). For two spike trains of units *x* and *y*, TE can be estimated as

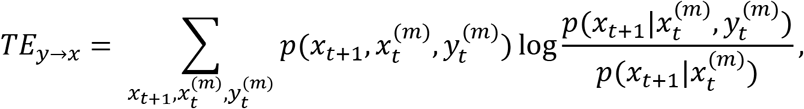

where *m* denotes embedding dimension (pattern size), and p(∙) implies probability. 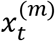 denotes *m* size spike pattern. For example, for *m* = 3 cases, there are 8 (=2^3^) possible spike patterns ([0,0,0], [0,0,1], …, [1,1,1]). TE_*y*→*x*_ (TE_*x*→*y*_) measures the statistical influence of unit *y* (*x*) on unit *x* (*y*). TE_*y*→*x*_ is the reduced amount of uncertainty in future of *x* by knowing the past of *y* given past of *x*. TE can also be thought of a mutual information (*I*) between *x* and 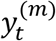 given past of 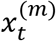 :

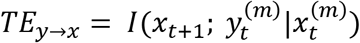

We used *m* = 3, and all spike trains were down-sampled to 125 Hz before calculation of TE; that is, the individual value in each bin of the spike train is one if there is one or more spikes within the 8 ms bin and zero otherwise.

### Multi-unit spike properties

Three measures, the total number of spikes (TNS), longest period below mean (LPBM), and sample entropy (SpEn), were calculated with MUA signal (i.e., sum of all SUAs). TNS represents the amount of total spike events occur in a sampled neural network. For the calculation of LPBM and SpEn, the MUA signal was convolved with Gaussian kernel with standard deviation of 25 ms (Vyazovskiy et al., 2011), and continuous spike signal was obtained. Details of LPBM and SpEn estimation are described below.

*Longest period below mean (LPBM)* LPBM measures the time length of the longest inactive periods (or active periods depending on the time series characteristics) in a given time series (Lubba et al., 2019); first, the time lengths of all consecutive values below the mean of time series are calculated and then the maximum of the time lengths are obtained as a LPBM value. LPBM is known to be one of the important temporal statistics in time series analysis (Lubba et al., 2019), and was used in this study to be a measure of persistent inactiveness of spike activity. A high LPBM value in MUA signal implies an existence of long inactive period suggesting synchronous fragmentation of spike activities, whereas a low LPBM indicates more continuous activity suggesting an irregular and asynchronous spiking pattern. Therefore, LPBM is expected to yield a high value when spikes are synchronously fragmented in time (e.g., slow oscillation and burst-suppression). In addition, LPBM will further increase as burst-suppression ratio increases with deepening of anesthesia.

*Sample entropy (SpEn)* SpEn was used to estimate the statistical irregularity of MUA as a time series. SpEn is an approximation of Kolmogorov entropy that measures the predictability of consecutive time series values based on their past values (Richman et al., 2000). A high SpEn value implies random or unpredictable dynamics while a low SpEn value indicates regular or deterministic dynamics. SpEn has been used to quantify depth of anesthesia and the level of consciousness in EEG studies (Liang et al., 2015; Liu et al., 2018), and it generally decreases as anesthetic deepens. To calculate the SpEn, first an embedded time series is obtained,

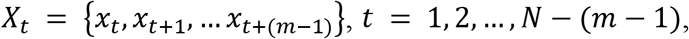

where *x*_*t*_ is time series value (convolved MUA signal in this study) at time*t*, and *m* is embedding dimension (pattern size). Second, the correlation sum is calculated from the embedded time series,

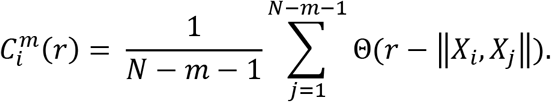

where Θ(∙) denotes a Heaviside step function and ‖∙‖ implies Euclidean distance between two vectors, and *r* represents the distance criteria. We used *m* = 3, and *r* = 0.2 standard deviation of amplitudes within each epoch following previous literature (Liang et al., 2015; Liu et al., 2018). Finally, the SpEn is defined as,

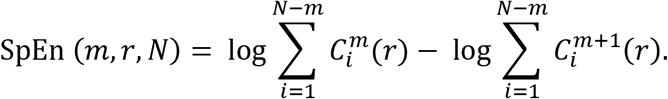

Before the SpEn calculation, the MUA signal was convolved as in the case of LPBM calculation, and down-sampled to 125 Hz.

### Classification of brain states

The primary focus of the study was to examine brain state-dependent changes in spike activity patterns during and after anesthesia. To this end, spike train data was first segmented into 10 second nonoverlapping epochs. Then five features from population level spike activity, that is, the total number of spikes (TNS), mean of log-transformed spike rate (SR_m_), mean local variation (LV_m_), longest period below mean (LPBM), and sample entropy (SpEn) were measured in each epoch. The five features have different ranges with each other, and different animals often show different ranges for a single feature.

Therefore, to normalize the feature values and mitigate the effect of outliers, the feature data were divided into sextiles in each animal, and were transformed by linearly scaling to a given range (0-1); that is, the median of the data in the first sextile was considered zero and the median of data in the last sextile was considered one in each experiment. This procedure is based on an assumption that the range of each of the five features is similar across different experiments with the same anesthetic protocol.

The five features were then used for unsupervised clustering to delineate distinct brain states. Hierarchical agglomerative algorithm with Ward’s linkage method were applied for the clustering of brain states, using Python package Scikit-Learn (www.scikit-learn.org). Each data point of a 10 second epoch was first treated as a single cluster in feature space, then the points were successively merged until all clusters merged into a single cluster. The method does not require a specific number of clusters at the beginning step, and the clusters can be easily identified from the hierarchy tree (dendrogram) that is built from the algorithm. We determined the optimal number of clusters based on the dendrogram and so-called elbow method. A within-cluster distance was plotted against the number of clusters, and the point where the curve sharply bends was chosen as an “elbow” point. We used the maximum of the 2^nd^ order difference of the distance-*K* curve to find the elbow point. We neglected the *K* = 2 cases, in which the brain state simply represents anesthetized (6-2% desflurane) and waking state (0% desflurane). In our preliminary studies, adding more features and performing principal component analysis barely changed the clustering results.

### Statistical analysis

All statistical analyses were conducted using StatsModels library (www.statsmodels.org) in Python 3.7. For all measures, to test the difference across the brain states, statistical comparisons were first performed using linear mixed models (LMM) based on restricted maximum likelihood estimation. For all LMMs, the brain states (categorical independent variable) were used as a fixed effect. For the properties of population activity (i.e., PSD of LFP, the five input features, and EMG), the random effect included the seven animals. For the individual unit properties such as SR and LV, the random effect included the different animals and units. Post-hoc pairwise comparisons were made between the brain states using a Bonferroni adjusted p-value < 0.05 (number of hypotheses = 10).

## RESULTS

### Cross-correlogram between SUA and MUA

The classification of putative excitatory (pE) and inhibitory (pI) units was conducted based on spike waveform and autocorrelogram as described in method section. 36 out of 251 units were classified as pI unit (14.4%). We further confirmed the classification by examining the asymmetry of cross-correlogram between SUA and MUA. As predicted, the asymmetry of pI (pE) units showed negative (positive) values on average (Fig. 2E); statistical significance was seen both in pE and pI units (one sided *t*-test with Bonferroni correction, p = 0.045 for pE, and p < 10^−6^ for pI). This suggests that pI (pE) units on average, tend to inhibit (promote) population activity, reassuring the classification of neuronal types.

### Brain state shifts during anesthesia

In order to identify local brain states from the electrophysiological recording independent of the nominal anesthetic concentration, we visualized how LFP spectrogram and MUA characteristics change over time during the experiment. In all animals, the LFP spectrogram, total number of spikes (TNS), mean of log-transformed spike rate (SR_m_), mean of local variation (LV_m_), longest period below mean (LPBM), and sample entropy (SpEn) profoundly changed during (6,4, and 2% desflurane) and after (0% desflurane) anesthesia. Figure 2A illustrates the time course of these variables in one animal as an example. Importantly, both the spectrogram and the MUA features change not only between but also within each recording period at constant anesthetic concentration. For instance, in the middle of the recording at 6% desflurane, low frequency (< 4 Hz) power in the LFP spectrogram and LPBM abruptly increased for no evident reason. The additional abrupt transitions are seen at 2% desflurane. Other animals also showed similar transitions, with either positive or negative sign, at various anesthetic levels (data not shown). This example demonstrates that a simple one-to-one relationship between the chosen LFP/MUA variables and the anesthetic concentration does not exist suggesting the need for a more nuanced identification of brain states from these variables.

To achieve this goal, we used agglomerative clustering on the five MUA variables as input features from data pooled from all animals to identify distinct, unitary brain states. The scatter plots in Fig. 2B illustrate pairwise relationships of the five MUA features in 5 clusters. The choice of 5 clusters could be justified by the dendrogram (Fig. 2C), which illustrates that between-cluster distances were large and within-cluster distances were small at *K* = 5. We also calculated a within-cluster distance as a function of *K* (Fig. 2D). The 2^nd^ order difference of the distance curve was maximized at *K* = 5, suggesting it was an optimal choice consistent with the dendrogram distances.

The five clusters identified by unsupervised clustering were designated as brain states S1 to S5 and the mean values of MUA variables among these states were statistically compared (Fig. 2E; Table 1). As found, S1 was characterized by the lowest TNS, SR_m_, and SpEn and the highest LPBM indicating that S1 corresponded to burst-suppression (see Fig. 3A) typical to deep anesthesia. In fact, S1 was mostly observed at 6% desflurane (Fig. 2F-G). S5, on the other hand, was mostly observed at 0% desflurane (Fig. 2F). It was characterized by high spike activity (high TNS and SR_m_) and asynchronous firing patterns (high SpEn and low LV_m_). S2 and S4 had intermediate feature values between those of S1 and S5. S2 was mostly observed at 4% desflurane and S4 was mostly seen at 2% desflurane (Fig. 2F).

**Table 1.** P-values of post hoc test for the five features. These features were used as an input to the clustering algorithm for the brain state classification. P-values were Bonferroni corrected.

|  | S1-S2 | S1-S3 | S1-S4 | S1-S5 | S2-S3 | S2-S4 | S2-S5 | S3-S4 | S3-S5 | S4-S5 |
| --- | --- | --- | --- | --- | --- | --- | --- | --- | --- | --- |
| TNS | $< 10^{-9}$ | $< 10^{-25}$ | $< 10^{-36}$ | $< 10^{-99}$ | $< 10^{-3}$ | $< 10^{-8}$ | $< 10^{-46}$ | 0.295 | $< 10^{-24}$ | $< 10^{-15}$ |
| SR<sub>m</sub> | $< 10^{-3}$ | 0.578 | $< 10^{-22}$ | $< 10^{-56}$ | 0.135 | $< 10^{-6}$ | $< 10^{-30}$ | $< 10^{-14}$ | $< 10^{-44}$ | $< 10^{-7}$ |
| LV<sub>m</sub> | 2.621 | $< 10^{-18}$ | $< 10^{-13}$ | $< 10^{-61}$ | $< 10^{-13}$ | $< 10^{-9}$ | $< 10^{-53}$ | 1.900 | $< 10^{-12}$ | $< 10^{-17}$ |
| LPBM | $< 10^{-19}$ | $< 10^{-59}$ | $< 10^{-26}$ | $< 10^{-37}$ | $< 10^{-11}$ | 0.750 | 0.002 | $< 10^{-6}$ | 0.007 | 0.451 |
| SpEn | $< 10^{-3}$ | $< 10^{-55}$ | $< 10^{-25}$ | $< 10^{-76}$ | $< 10^{-30}$ | $< 10^{-9}$ | $< 10^{-45}$ | $< 10^{-5}$ | 0.068 | $< 10^{-13}$ |

**Figure 3.**
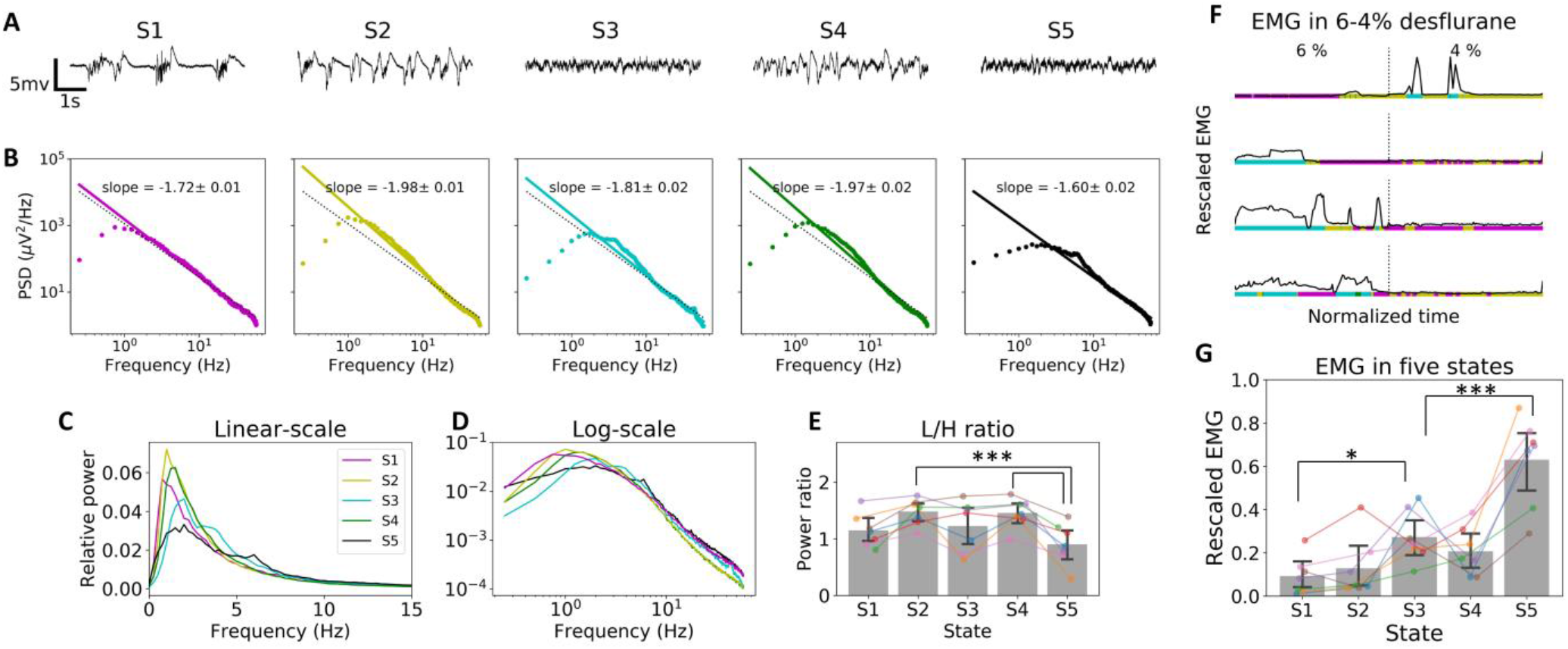
Local field potential (LFP) and electromyography (EMG) in five brain states. (A) Example LFP traces from one animal. (B) Power spectral density (PSD) vs. frequency plot shows power-law relationship in the five brain states. Data were averaged from the seven animals. (C) Relative PSD vs. frequency plot in a linear scale. (D) Log-scale representation of the PSD-frequency plot. (E) L/H power ratio across five brain states. The L/H ratio is defined as log10{(PSD at 0.25-4 Hz)/ (PSD at 30-59 Hz)}. Error bar indicates 95% confidence interval across animals (***p<0.001 compared to S5). (F) EMG activity (black trace) during 6-4% desflurane in four animals. Horizontal bars with different colors indicate different brain states; magenta, yellow, cyan, and green for S1, S2, S3, and S4, respectively. (G) EMG in S3 is significantly higher than that of S1 and lower than that of S5. Error bar indicates 95% confidence interval across animals (*p<0.05, ***p<0.001 compared to S3).

Interestingly, S3 was mostly found in 6% desflurane (Fig. 2F) similar to S1. However, S3 showed a distinct pattern from S1. It was characterized by high SpEn, relatively low SR_m_ and very low LBPM. TNS was not reduced as much as SR_m_; Notice that TNS indicates total number of spikes in the neuronal population and SR_m_ is the mean of log-transformed individual spike rates. The discrepancy suggests that in S3, some neurons are inactive, but a few neurons emit a large number of spikes whereas these outliers are absent in S1. The details of spike rate distribution of neuronal population are analyzed further in the next section. The low LPBM indicates no clear distinction of active and inactive period in S3. The low LPBM, together with the high SpEn, implies that spiking pattern in the S3 was asynchronous.

Although brain state and anesthetic concentration were not uniquely related, a general trend of the occurrence probability of brain states with anesthetic level was evident; the S1, S2, S4, and S5 in order occurred mostly at correspondingly decreasing desflurane concentration. Therefore, it is reasonable to surmise that the occurrence probability of brain states, with an exception of S3, generally reflected the depth of anesthesia. Interestingly, however, they were also observed in other concentrations (Fig. 2G-F). For instance, at 6% desflurane, S1 was present 61% of the time, whereas at 4% desflurane, S1 was present 39% of the time; with the balance occupied by other brain states. Generally, several different brain states occurred at each constant anesthetic concentration. For example, at 6% desflurane, S1, S2, S3, and S4 appeared in non-negligible proportion (Fig. 2G). The many-to-many relationship between brain state and anesthetic concentration suggests a general need for brain state-dependent investigation of unit activity.

### LFP properties of the five brain states

Because local field potentials (LFP) generally reflect the state in anesthesia, we examined LFP patterns and power spectral density (PSD) in each brain state. Typical LFP traces in five brain states are shown in Figure 3A from the same animal as in Fig. 2A. The LFP in S1 exhibited burst-suppression. S2 and S4 revealed relatively high amplitude, slow activity as generally expected in anesthesia. In contrast, S3 showed low amplitude, desynchronized LFP pattern similar to S5 corresponding to the awake state. The PSD averaged over all animals showed a power law relationship with frequency (Fig. 3B). The slightly higher slope in S2 and S4 was associated with the increased low-frequency (< 4 Hz) and decreased high-frequency (> 30 Hz) PSD (Fig. 3C) consistent with the anesthetic-induced suppression of high frequency gamma power and enhancement of delta and slow oscillation in EEG/LFP. S5 was characterized by increased theta (5-9 Hz) and high-frequency (> 20 Hz) power, the typical signatures of EEG/LFP in wakefulness. For a quantitative comparison we calculated the L/H ratio as log10{(PSD at 0.25-4 Hz)/ (PSD at 30-59 Hz)} (Li et al., 2009). S2 and S4 showed significantly higher L/H ratio than S5 (p<0.001; Fig. 3E). In sum, the brain states, S1, S2, S4 are consistent with known LFP features of deep, moderate and light anesthesia, respectively; however, the LFP in S3 is unexpected and contrary to the generally presumed dose-dependent effect of anesthesia.

### High EMG activity in paradoxical desynchronized state

The asynchronous firing pattern and relatively high LFP gamma power found in S3 raises the question whether the systemic arousal level may also be elevated in S3 as it is in S5. Generally, the EMG follows the level of arousal; therefore, the vigilance state of animals was estimated by EMG activity. Although both S1 and S3 occurred mostly in 6% desflurane, EMG of S3 was substantially higher than that of S1. The rescaled EMG traces from each animal exhibited higher muscle activity in S3 than in S1 and sometimes even higher than in S2 (Fig. 3F). Statistically significant differences in the rescaled EMG were found for S3 vs. S1 and S3 vs. S5 (p < 0.024 and p < 10^−6^, respectively with Bonferroni correction; Fig. 3G).

### Spike rate distribution across brain states and neuron types

We compared the five brain states in terms of both the number of emitted spikes and the average spike rate of individual units. Generally, desflurane suppressed spike activity (Fig 4A-B). Figure 4A illustrates the time course of SR_m_ (average of log-transformed spike rate) and total spike number TNS from the same animal as in Figure 2A and 3A. As seen there, the traces of SR_m_ and TNS deviated from each other, especially at 6% desflurane. The TNS showed a pronounced decrease when the brain state transitioned from S3 to S1, whereas the SR_m_ remained the same. In S3 many units were inactive, even more than in S1 and S2, but there were a few units with very high SR (Fig. 4C). Accordingly, the variation of SR across individual units was the highest in S3. The variation in SR distribution was quantified by the Gini coefficient and the value of S3 was significantly larger than all others (Fig. 4D; Table 2). Specifically, Figure 4E implies that the highly active units in S3 are putative excitatory (pE) units. SR of active units in S3 was comparable to that in S5 (left panel in Fig. 4E) for pE units; however, SR of putative inhibitory (pI) active units in S3 was lower than that of pI units in S5 (right panel in Fig. 4E). For the mean SR of pE units, there were significant differences among the states except S1 vs. S3; SR_m_ increased from S1 or S3 through S2 and S4 to S5 (Fig. 4E; Table 2). For pI units, SR_m_ was significantly higher in S5 than in all other states. Thus, in general, desflurane profoundly suppressed SR of both pE and pI units, but a few pE units in S3 remained highly active resulting in very high SR variation in this brain state.

**Table 2.** P-values of post hoc test for all SUA features. P-values were Bonferroni corrected.

|  | S1-S2 | S1-S3 | S1-S4 | S1-S5 | S2-S3 | S2-S4 | S2-S5 | S3-S4 | S3-S5 | S4-S5 |
| --- | --- | --- | --- | --- | --- | --- | --- | --- | --- | --- |
| SR (pE) | $< 10^{-6}$ | 1.164 | $< 10^{-30}$ | $< 10^{-81}$ | $< 10^{-3}$ | $< 10^{-8}$ | $< 10^{-40}$ | $< 10^{-23}$ | $< 10^{-69}$ | $< 10^{-12}$ |
| SR (pI) | 0.875 | 8.112 | $< 10^{-3}$ | $< 10^{-16}$ | 1.416 | 0.108 | $< 10^{-10}$ | $< 10^{-3}$ | $< 10^{-15}$ | $< 10^{-3}$ |
| Gini coef. | 5.108 | $< 10^{-3}$ | 2.526 | 1.433 | $< 10^{-4}$ | 6.267 | 4.202 | $< 10^{-5}$ | $< 10^{-6}$ | 7.493 |
| BR (pE) | 0.001 | $< 10^{-13}$ | $< 10^{-8}$ | $< 10^{-26}$ | $< 10^{-3}$ | 0.089 | $< 10^{-12}$ | 0.695 | 0.013 | $< 10^{-6}$ |
| BR (pI) | 1.170 | 0.003 | 2.359 | 4.400 | 0.260 | 6.570 | 3.690 | 0.055 | 0.012 | 6.339 |
| LV (pE) | 0.350 | $< 10^{-36}$ | $< 10^{-22}$ | $< 10^{-114}$ | $< 10^{-30}$ | $< 10^{-16}$ | $< 10^{-111}$ | 0.003 | $< 10^{-22}$ | $< 10^{-47}$ |
| LV (pI) | 5.354 | 0.157 | $< 10^{-6}$ | $< 10^{-13}$ | 0.013 | $< 10^{-9}$ | $< 10^{-18}$ | 0.014 | $< 10^{-7}$ | 0.046 |
| Corr (pE) | $< 10^{-20}$ | $< 10^{-159}$ | $< 10^{-96}$ | $< 10^{-243}$ | $< 10^{-77}$ | $< 10^{-32}$ | $< 10^{-144}$ | $< 10^{-12}$ | $< 10^{-7}$ | $< 10^{-45}$ |
| Corr (pI) | 0.006 | $< 10^{-35}$ | $< 10^{-8}$ | $< 10^{-40}$ | $< 10^{-21}$ | 0.018 | $< 10^{-25}$ | $< 10^{-11}$ | 7.046 | $< 10^{-14}$ |
| TE<sub>s</sub> | 1.132 | 8.272 | 0.003 | $< 10^{-12}$ | 0.715 | 0.434 | $< 10^{-7}$ | 0.001 | $< 10^{-13}$ | 0.001 |
| TE<sub>m</sub> | 0.059 | $< 10^{-4}$ | 0.001 | $< 10^{-6}$ | 0.457 | 1.505 | 0.075 | 5.753 | 4.999 | 2.169 |
| TE<sub>m</sub> (pE-to-pE) | 0.017 | $< 10^{-4}$ | $< 10^{-5}$ | $< 10^{-8}$ | 1.633 | 0.334 | 0.016 | 4.639 | 0.775 | 3.018 |
| TE<sub>m</sub> (pI-to-pE) | 0.002 | $< 10^{-8}$ | 0.001 | $< 10^{-5}$ | 0.094 | 8.457 | 1.205 | 0.163 | 2.966 | 1.744 |
| TE<sub>m</sub> (pE-to-pI) | 0.102 | $< 10^{-5}$ | 0.094 | 0.031 | 0.115 | 9.764 | 6.852 | 0.125 | 0.338 | 7.07 |
| TE<sub>m</sub> (pI-to-pI) | 3.434 | 0.071 | 5.831 | 3.411 | 0.645 | 1.126 | 9.968 | 0.006 | 0.604 | 1.086 |
| Gini coef. (TE) | 2.576 | 2.462 | 0.002 | $< 10^{-13}$ | 0.219 | 0.093 | $< 10^{-9}$ | $< 10^{-5}$ | $< 10^{-17}$ | 0.001 |

**Figure 4.**
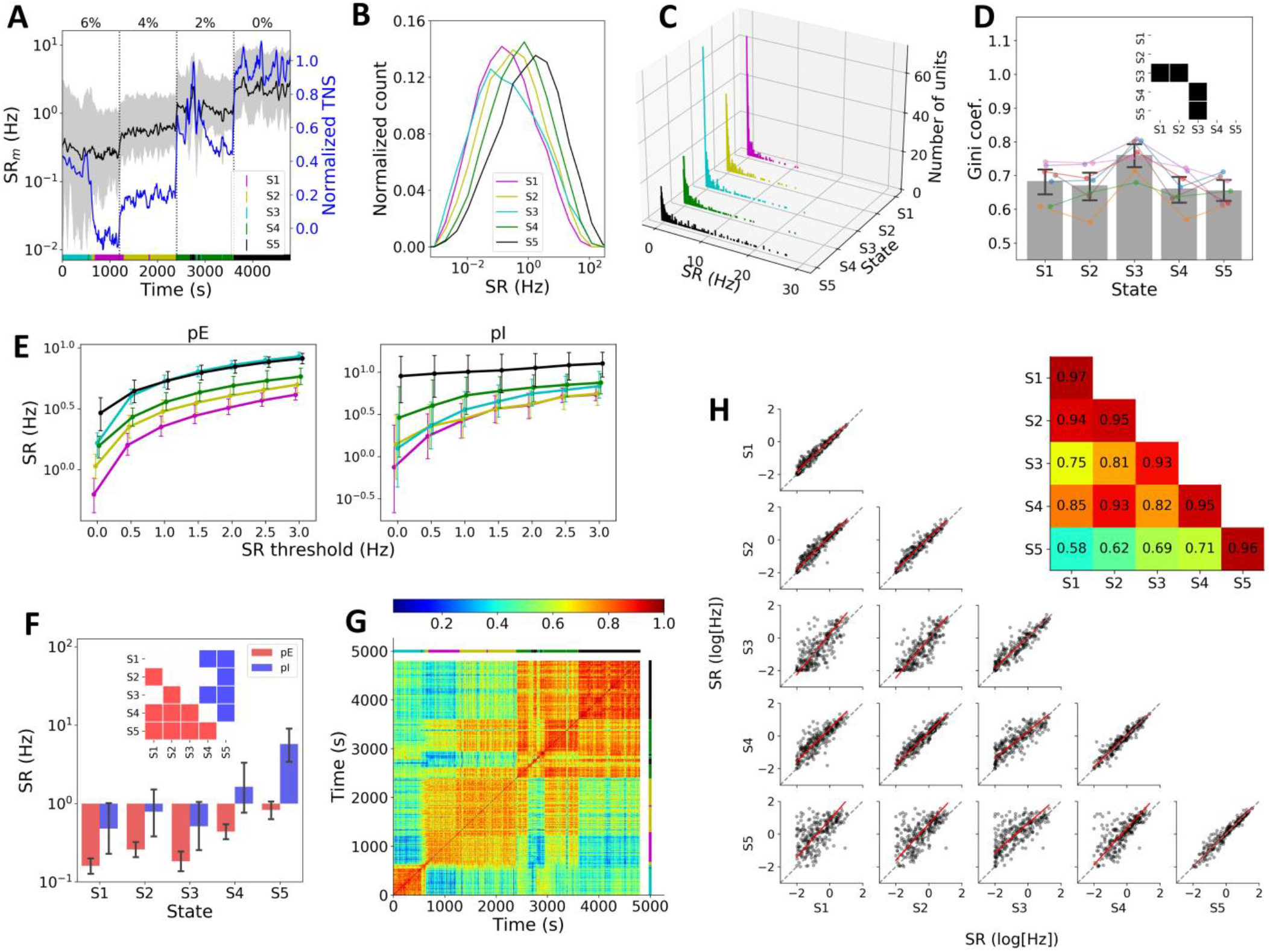
Spike rate properties of five brain states. (A) Example trace of mean spike rate (SR_m_, black) and total number of spikes (TNS, blue) from one animal. SR_m_ is the average of log-transformed spike rates. Black shaded area represents 25th and 75th percentiles of log-transformed spike rate distribution. The horizontal bars with different colors indicate different brain states. (B) Log-transformed spike rate distribution in five brain states from all individual units. (C) Frequency distribution of untransformed spike rate (SR) in five states. Color code is the same as in panel A and B. (D) Gini coefficient of untransformed spike rate distribution. Error bar indicates 95% confidence interval across units. The inset represents pairwise statistical significance. (E) Average log-transformed spike rate as a function of SR threshold for pE (left) and pI (right) units. (F) Comparison of SR_m_ across five states. The inset shows statistically significant difference for pairs of brain state. (G) Pairwise spike rate correlation among all epochs from the same animal in panel A. Two color bars above and right of the matrix indicate the five brain states (same color code as before). (H) Orthogonal linear regression of log-transformed spike rates between brain states. The inset in the upper right corner represents Pearson correlation values for within- and between-brain states.

A decrease in average firing rate could be generalized across all units or selective to specific units; e.g. due to a slowing of high-firing neurons. Therefore, we tested if units had a tendency to preserve their firing rate rank across brain states. SR similarity between any two epochs was estimated by calculating the Pearson correlation, and is presented in Figure 4G. The correlation matrix for individual units showed high within-state similarity and relatively low between-state similarity. The results from correlation analysis of all units from all animals (n = 251; Fig. 4G) were consistent with the result from the representative animal. Within-state comparisons of SR between the first half of the state and the second half of the same state are shown in the diagonal panels of Fig. 4G. Between-state comparisons between different states are shown in the off-diagonal panels of Fig. 4G). Orthogonal linear regression indicated that within-state similarity of SR (R > 0.93 for all the five states) is generally higher although still significant (p < 0.001) than between-state similarity except S1 vs. S2 (upper right inset in Fig. 4G). These findings indicate that SR profiles of individual units are preserved both within and between brain states.

### Temporal dynamics of spike activity

Anesthesia not only suppresses the average spike rate as reported in the previous section (Fig. 4) but also modulates the temporal dynamics of spike activity (Vizuete et al., 2014). Raster plots in Figure 5A illustrate the changing temporal dynamics of spike activity. Note that in S1, S2, and S4, but not in S3, spike activity is more synchronized and temporally fragmented as compared to S5. Figure 5B displays raster and distribution of ISIs in seven representative units (four pE and three pI) from the same animal at different desflurane concentrations. The shape of ISI distribution was profoundly altered by the anesthetic. In wakefulness (S5) the ISI distribution was unimodal, whereas in the other four states it was bimodal or multimodal. This was partially due to the silent periods in spike activity; the large ISI values in the raster plot (ISI ~ 10^3^ ms) in Figure 5B (especially, S1 and S2) correspond to silent periods that contribute to a second peak in ISI histogram (Fig. 5B). In addition, some pE units in S1 and S2 tended to fire in brief bursts that were associated with short ISI (ISI < 10^1^ ms; Fig. 5B). Burst activity had also contributed to a peak near ISI ~ 10 ms in the ISI histograms (Fig. 5B). In S3, two pE units exhibited very high SR (represented by dense points, asterisk in Fig. 5B) that was comparable to the SR in S5, consistent with the findings in the previous section (Fig. 4E). Both units showed unimodal ISI distribution as in S5.

**Figure 5.**
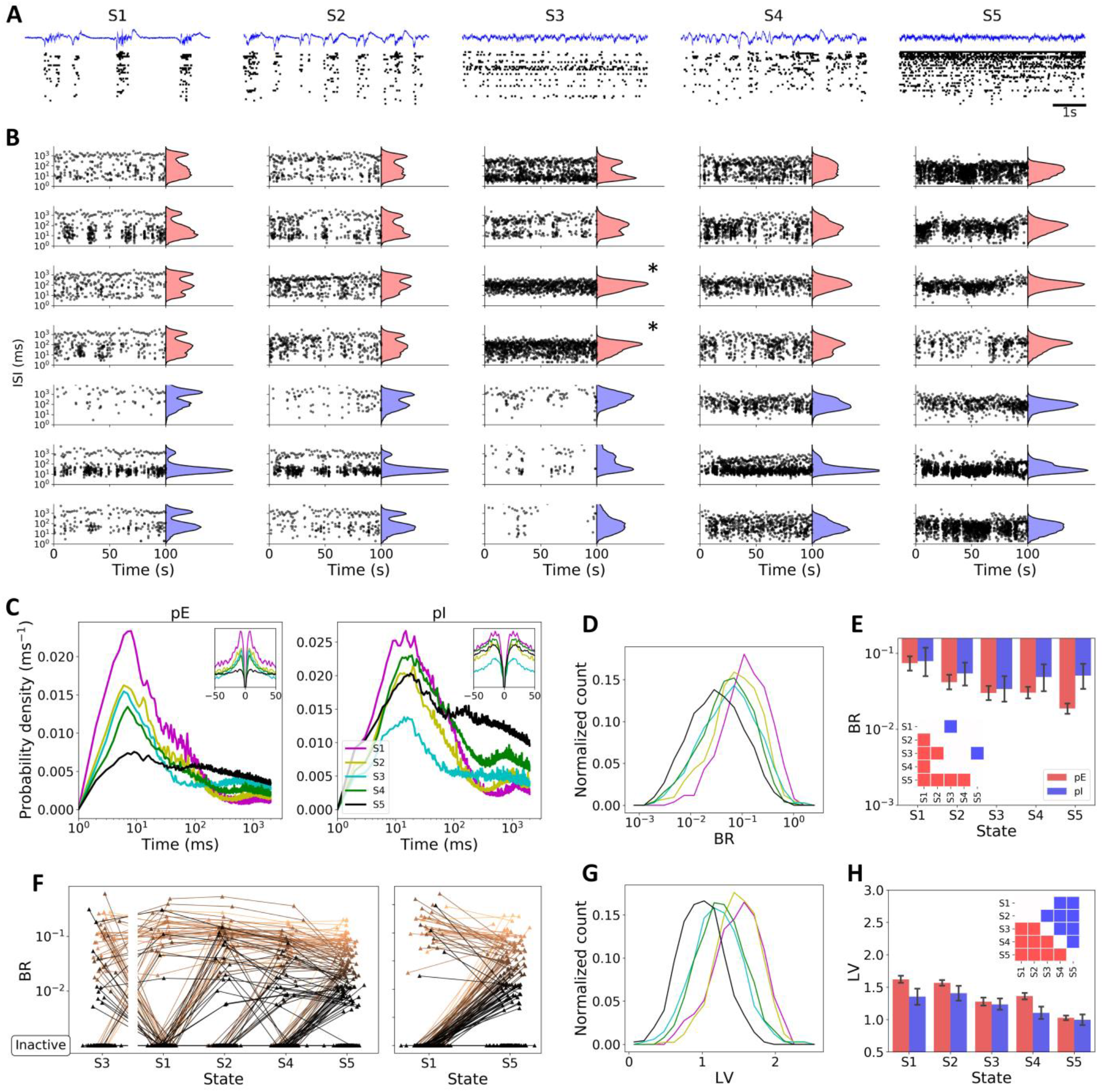
Temporal dynamics of individual units. (A) Example local field potential trace and corresponding raster plot of single units in one animal in brain states S1 through S5. (B) Raster plot and frequency distribution of inter-spike interval (ISI) of seven units from the same animal in panel A; red histograms: pE units, blue histograms: pI units. (C) Average normalized autocorrelogram on log-scale calculated from all active units (SR≥1 Hz). Inset: the same autocorrelogram on linear scale for short time lags (−50 to 50-ms). (D) Distribution of burst ratio (BR) of active units (SR>1 Hz). (E) Comparison of BR across five states. Error bar indicates 95% confidence interval across units. The inset shows statistically significant pairs of brain state. (F) An illustration of the change of BR and in(activeness) with brain state in individual pE units shows that pE units become either bursty or inactive under anesthesia; each line connects data from the same unit. A dark (light) colored line indicates low (high) BR value in S5. Inactive units (SR<1 Hz) are shown at the bottom of each panel. For better visualization of the gradual change from S1 through S2 and S4 to S5, data in S3 were separated to the left. The right panel emphasizes two extreme cases (S1 and S5). (G) Distribution of local variation (LV) of active units (SR<1 Hz) only. (H) Comparison of LV across five states. Error bar indicates 95% confidence interval across units. The inset represents statistically significant difference for pairs of brain state.

To determine if burst activity and long silence periods generally occurred in all units and all animals, autocorrelogram was calculated from all active units (SR ≥ 1 Hz). The averaged autocorrelogram showed a gradual increase of burst activity (ISI < 10 ms) in pE units under anesthesia from S5 to S1 (left panel in Fig. 5C). A prominent peak was observed near at 6 ms that progressively decreased from S1 to S5 (left panel in Fig. 5C). As expected, autocorrelogram showed little or no evidence of bursting of pI units. Another measure of bursting of pE units, the burst ratio (BR) generally decreased from S1 to S5 (Fig. 5D-E). Note that inactive units (SR < 1 Hz) were excluded from the autocorrelogram and BR calculation. Similar to the SR distribution, BR did not follow normal distribution but skewed to the right, and thus it was log-transformed. Statistically significant difference in BR of pE units was found for all pairwise comparisons of brain states except S4 vs. S2, and S3 (Fig. 5E; Table 2). The increases in BR of pI units were less pronounced (right panel in Fig. 5C; Fig. 5E). For pI, BR of S3 was significantly lower than that of S1 and S5. The suppression in SR (Fig. 4), together with the changing temporal pattern of spiking indicates that neurons were either inactive or bursty at deeper levels of anesthesia. As the brain state changed from S1 to S5, more units became active and burst activity of active units in anesthesia was reduced in S5 (Fig. 5F). Again, S3 was an exception; BR of pE units in S3 was comparable to BR in S4.

The state-dependent changes of ISI distribution were also characterized by local variation (LV), a measure of spike timing variability - a measure that is sensitive to changes in both burst activity (small ISI) and to the presence of long silent periods (large ISI). LV showed a similar trend to BR across brain states but more effectively distinguished the five brain states especially for pI units (inset in Fig. 5E, H). LV of both pE and pI units decreased from S1 through S2 and S4 to S5. All pairwise comparisons except S1 vs. S2 were statistically significant for pE units and all pairwise comparisons except S1 vs. S2, and S3 were significant for pI units (Fig. 5H; Table 2).

In summary, spiking activity of most units was profoundly inhibited by desflurane and the remaining active units showed an enhanced burst activity (for pE) and prolonged silence period (both pE and pI). In the paradoxical state S3 these effects were marginal such that units exhibited an irregular spiking pattern similar to that seen in wakefulness or S5.

### Individual neurons conform to population activity

It is well known that desflurane, as well as other anesthetics, enhances spike-field correlation (Vizuete et al., 2014). We re-examined this effect as a function of brain state and found that desflurane increases spike-LFP correlation in all anesthetized states except S3. Specifically, the spike-triggered LFP amplitude decreased from S1 through S2, and S4 to S5 but not in S3 (Fig. 6A). We also examined spike-triggered MUA (Fig. 6B). The MUA in S5 was high and relatively flat across time lags, with a small oscillatory pattern in theta frequency range (5-9 Hz). From S4 through S2 to S1, the overall MUA level gradually decreased indicating a suppression of overall spike activity, while the MUA peak near 0 ms remained almost the same indicating synchronous firing. In addition, the “dip” in MUA observed before and after spike events became deeper and wider as brain state moved from S4 through S2 to S1. Notice that in S1, the number of spikes near ± 500 ms to spike events is close to zero, consistent with the near-silent periods of LFP burst-suppression. Again, distinct from the other three anesthetic states, S3 did not have a large trough on either side of spike events; whereas the MUA was substantially lower than in S5.

**Figure 6.**
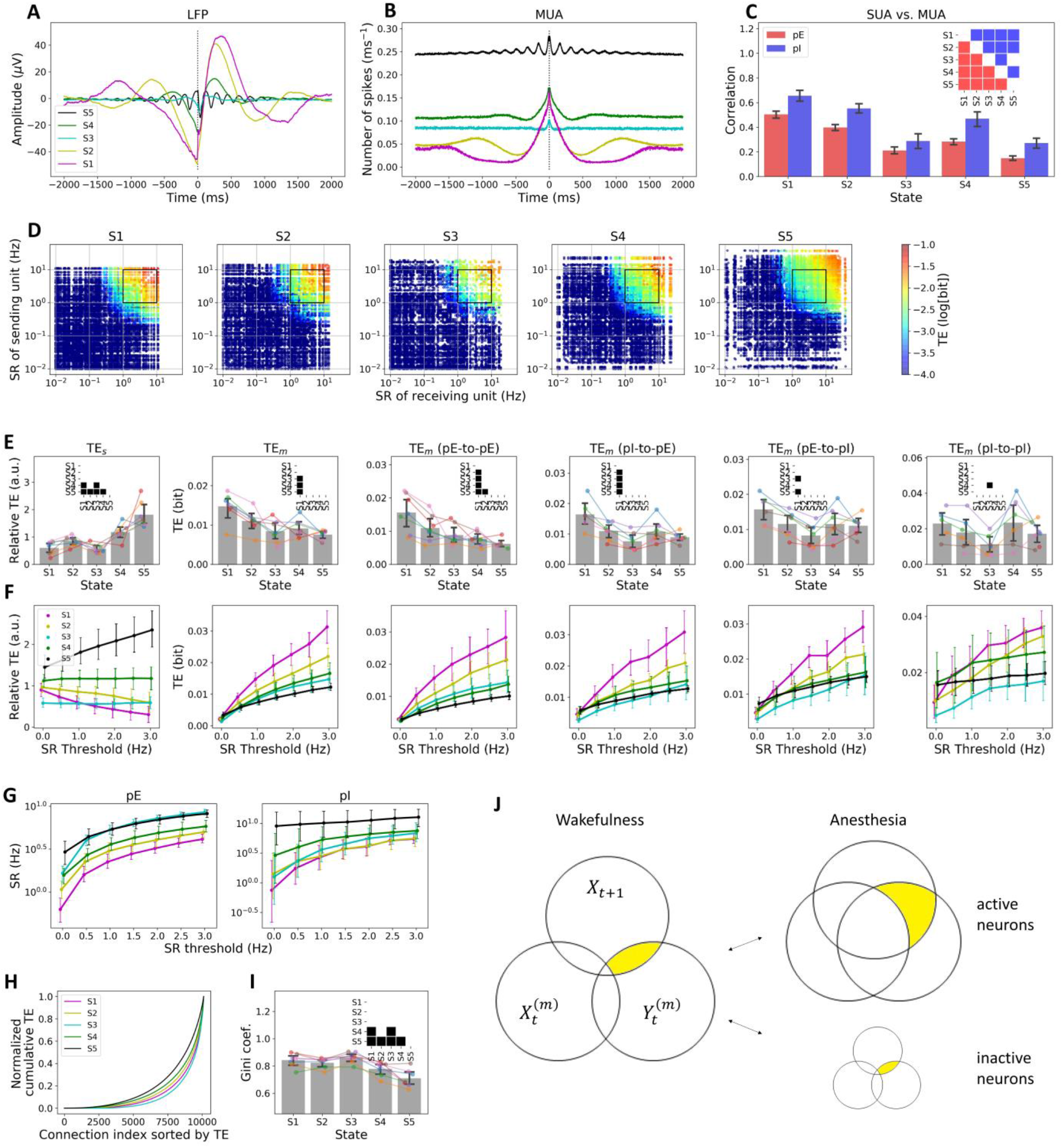
Functional connectivity in five brain states. (A) Spike-triggered average local field potential (LFP) in brain states S1 through S5. (B) Same for spike-triggered average multi-unit activity (MUA). (C) Pearson correlation between single-unit activity (SUA) and MUA. Error bar indicates 95% confidence interval across units. The inset represents statistically significant difference for pairs of brain state. (D) All pairwise transfer entropy (TE) of individual units from all animals on the axis of spike rate of individual units. The color of each dot in each panel represents TE value between two units (red for high TE, blue for low TE). The x-axis (y-axis) corresponds to spike rate of information-receiving (-sending) unit. (E) Comparison of TE of active units (SR≥1 Hz) across five brain states. Error bar indicates 95% confidence interval across animals. The inset represents statistically significant difference between brain states. (F) TE as a function of spike rate threshold; i.e., TE values of active units are averaged while criteria for active and inactive units varies. States correspond to those as in the panel above. (G) Average log-transformed spike rate pf pE and pI units as a function of spike rate threshold. (H) Lorenz curve of pairwise TE (including inactive units) from pooled data. (I) Gini coefficient of the TE distribution. Error bar indicates 95% confidence interval across animals. The inset represents statistically significant pairs of brain state. (J) Information Venn diagram to illustrate the relationship between two neurons, *X* and *Y* in wakefulness and anesthesia. The yellow area indicates the amount of information transfer from *Y* to *X* (*TE*_*y*→*x*_). High spiking activity in wakefulness suggests high TE (left). In anesthesia, neurons are either inactive or intermittent/burst firing with enhanced synchronization. TE between inactive neurons is extremely small, whereas TE between active neurons is larger than the TE in wakefulness. This results in a large variation of TE values as seen in (H, I).

To evaluate the extent to which individual neurons conform to population activity, we quantified SUA-MUA correlation. Both spike train of individual single units and MUA signal were convolved with Gaussian kernel (SD = 25 ms), then the Pearson correlation between the two convolved signals was calculated. A substantial change in correlation was observed in both pE and pI units (Fig. 6C); all pairwise comparisons were statistically significant for pE units and all pairwise comparisons except S3 vs. S5 were significant for pI units (Table 2).

### Information transfer depends on spike rate

The correlation results described so far indicate a nondirectional relationship between convolved SUA and MUA signals. In order to estimate directional functional connectivity of neuronal interaction, TE between individual binary spike trains (SUAs) was calculated. Because spike activity itself is in part a result of neural communication, TE is presumed to depend on the degree of overall spike activity. In fact, TE was high for units with high SR and low for units with low SR (Fig. 6D). However, for a same range of SR, TE in anesthesia was higher than that in wakefulness. For example, for units having SR in a range of 10^0^ to 10^1^ Hz (the colored points inside the black squared box in each panel in Fig. 6D), TE values in S1 were higher than TE values in S5.

### Synchronous firing correlates with enhanced information transfer

The sum of TE values (TEs) for active units was highest in S5 and lowest in S1 and S3 (Fig. 6E left panel; Table 2), which is consistent with the general reduction of SR in anesthesia. However, the mean TE (TEm; the mean of TE values for active units) showed an opposite trend as it was the highest in S1 (Fig. 6E, second panel from the left; Table 2). Note that inactive units (SR < 1 Hz) were excluded from all results in Figure 6E and the averaging was done across animals (n = 7). To see if the SR exclusion threshold had any effect, TE results with different SR thresholds were compared in Figure 6F. When all units were included (SR threshold = 0 Hz), the TEm was the highest in S5 (Fig. 6F, second panel from the left) as it should be qualitatively the same as in TEs with SR threshold = 0 Hz (Fig. 6F, first panel from the left). However, as the SR threshold increased, TEm of S5 became the lowest and that of S1 the highest (Fig. 6F, second panel from the left). Figure 6G illustrates that this differences among brain states do not merely reflect the SR changes; that is, the mean SR in S5 was the highest of all states across all SR thresholds, distinct from the TEm for both pE and pI units. Note that for pE units, SR of S3 was comparable to that of S5 when SR threshold > 0 Hz.

The variation of all pairwise TE values (including inactive neurons) estimated by Gini coefficient was higher in anesthesia (S1-4) than wakefulness (S5) (Fig. 6H). This is because in anesthesia, many of the units were silent or inactive; thereby these units had very low TE, while the remaining active units had relatively high TE. Statistically significant differences in the Gini coefficient were found for S5 vs. S1, S2, S3, and S4 and S4 vs. S1, and S3 (Fig. 6I; Table 2).

The reason for the difference in change across brain states between the TEs and TEm can be attributed to the number of active neurons and (2) synchronous activity (Fig. 4C), and explained by Venn diagram of information in Figure 6J. In wakefulness, there are many active neurons; therefore, the sum of transfer entropies of active neurons (TEs) is high (left in Fig. 6J). In anesthesia, on the other hand, there are far less number of active neurons (Fig. 4B-C); therefore, the sum of transfer entropies of active neurons (TEs) is relatively small. For the small number of active neurons (upper right in Fig. 6J), the enhanced synchronization in anesthesia produces a large value of transfer entropy; this contributes the high value of the mean transfer entropy between active neurons (TEm). For inactive neurons, however, there is few information to be transferred, so that transfer entropy is extremely small (lower right in Fig. 6J). This also explains why the variation in transfer entropy is high in anesthesia (Fig. 6H-I).

### Information transfer along different connection types is state-dependent

We further investigated whether desflurane differentially affects different connection types, by examining TE of neurons pairs, pE-to-pE, pI-to-pE, pE-to-pI, and pI-to-pI (Fig. 6E-F). The most pronounced change with state was observed in pE-to-pE connectivity indicated by a gradual decrease of TE from S1 to S5 (Fig. 6E). Statistical significance was seen in S1 vs. all the other states and S2 vs. S5 (Table 2). The pI-to-pE connectivity was also higher in S1 vs. all the other states. The increase of TE in pE-to-pI and pI-to-pI connections was not as pronounced as in pE-to-pE and pI-to-pE cases. S3 showed relatively low TE such that pE-to-pI of S3 was lower than that of S1 (Fig. 6E, fifth panel from the left; Table 2) and pI-to-pI of S3 was lower than that of S4 (Fig. 6E, sixth panel from the left; Table 2). The findings suggest that desflurane exerts a more substantial effect on pE-to-pE and pI-to-pE connections than pE-to-pI and pI-to-pI connections.

## DISCUSSION

The main goal of this work was to determine how cortical unit activity changes with dynamically transitioning brain states under anesthesia. Using unsupervised machine learning method, we identified five brain states with distinct neuronal spiking behavior. Multiple brain states were observed at a constant anesthetic concentration, and conversely, the same brain state occurred at different anesthetic concentrations. The spontaneous shift of brain states at fixed anesthetic level suggested that the neuronal network underwent metastable (Bovier, 2006) or multistable state changes due to external perturbation or noise (Scott Kelso, 2012; Golos et al., 2015). Recent anesthesia studies of large-scale brain activity argued that neuronal dynamics may be at equilibrium on short timescales (seconds) but shows state switching at longer timescales (minutes) (Hudson et al., 2014; Hudson, 2017). Our results are consistent with these findings while providing additional insight into the spiking behavior of individual neurons in dynamically transitioning brain states.

### Spikes are synchronously fragmented in anesthesia

The intermittent firing pattern observed in anesthetized brain states (i.e., the increased LV and bimodal interspike intervals distribution) was synchronous among the neurons and therefore which was also reflected in the MUA, by an increase in LPBM. This synchronized fragmentation of spike activity estimated by the increase of individual-to-population correlation was more effective in distinguishing the four brain states (except S3) than all the other examined properties of spike dynamics suggesting that the synchronously fragmented spike activity is the most pronounced effect of anesthesia (Fig. 6C). A higher value of individual-to-population coupling implies that the spike activity of each neuron is constrained to the population activity. From the perspective of information processing, this must be an undesirable condition. Because the entire population acts like a single neuron, information capacity of the population is very low (Tononi, 2004; Izhikevich, 2006). Although high individual-to-population coupling suggests an increase of shared information among neurons, because information content of a single neuron is extremely limited, one could surmise that in such condition, the animal would be unconscious. This also explains why surgery is preferred in anesthetic states when EEG displays slow oscillations. Synchronized fragmentation of spike activity has also been reported with other anesthetics, such as ketamine/xylazine (Compte et al., 2003), urethane (Steriade et al., 1993; Kasanetz et al., 2002; Clement et al., 2008), propofol (Lewis et al., 2012), in addition to desflurane (Vizuete et al., 2014) despite the agents’ diverse molecular structure and pharmacological targets. Thus, our study suggests that synchronously fragmented spike pattern seen with most anesthetics is a common signature of impaired information processing closely associated with loss of consciousness.

### Unlike sleep, anesthesia may disrupt sensory functions

We found that desflurane reduced the spike rate of most neurons regardless of their wakeful firing rate unlike sleep that was found to differentially alter high-firing and low-firing neurons (Miyawaki and Diba, 2016). In natural sleep, the spike rate of high-firing neurons substantially decreased while the spike rate of low-firing neurons was enhanced (Watson et al., 2016). It has been suggested that high-firing neurons appear to be comprised of so-called *choristers*, which conform to the mean spike rate of the neuronal population, while the low-firing neurons called *soloists* respond to stimulation with firing rate changes distinct from that of the population (Bachatene et al., 2015). Specifically, the preferential augmentation of spike rate of low-firing, stimulus-selective neurons during rapid eye movement sleep has been thought to contribute to an increase of the signal-to-noise ratio of sensory processing. The fact that desflurane decreased the spike rate of virtually all neurons, including those with low baseline firing rate, may be one of the reasons why sensory functions fail in anesthesia.

### Anesthesia facilitates bursting of excitatory neurons

A recent modeling study of spiking neuronal network demonstrated that burst-spikes of individual neurons is more influenced by their presynaptic environment than by their cell type (Tomov et al., 2016). For example, regular spiking neurons could exhibit burst firing by network-mediated effect. Burst-spikes of individual neurons can also shape global network dynamics. In urethane anesthetized rats, burst-spikes induced by electrical stimulation of a single cortical neuron could switch global cortical state from slow oscillation (synchronized activity) to persistent UP state and vice versa (Li et al., 2009). Nevertheless, the causal relationship between the intrinsic spiking pattern of individual neurons and network synchronization has yet to be fully elucidated. While some neurons showed bursting, about three quarter of neurons were essentially inactive (fired at < 1 Hz) in deep anesthesia. Several modeling studies postulated the suppression of metabolic rate in the brain as a key mechanism of anesthetic-induced low frequency oscillations and burst-suppression (Cunningham et al., 2006; Ching et al., 2010, 2012). Another study reported the occurrence of burst-spikes is highly associated with suppression of spike activity, such that hyper-excitable state at the end of suppression period enables an emission of burst-spikes (Kroeger and Amzica, 2007). However, it remains uncertain how anesthesia almost completely suppresses majority of neurons while causing burst and synchronized activity in the remaining active neurons. Synchronous firing may be the only way for a neuronal network to maintain its activity under synaptic inhibition in anesthesia (Lukatch and MacIver, 1996); synchronous firing allows neurons to receive enough number of spikes from connected neurons within a short time period, thereby preventing from a decay of spike activity. In this scenario, neurons with many and strong synaptic connections would generate a large number of action potentials with high synchronization, and vice versa. According to recent study, neurons with strong population coupling (choristers) receive many synaptic inputs from their neighbors, and show high firing rate both in wakefulness and anesthesia (Okun et al., 2015). However, the existence of a paradoxical desynchronized state in deep anesthesia suggests the possibility of an alternative scenario in which neurons fire asynchronously while the mean firing rate is profoundly suppressed comparable to an averaged firing rate during burst-suppression period. Future modeling study of anesthesia which considers spike rate distribution and synaptic connections as well as the anesthetic-induced brain states may be able illuminate the possible causal relationship of spike rate distribution, burst activity, and synchronization.

### The paradoxical desynchronized state and consciousness

In the paradoxical desynchronized state (S3) which was mostly found during deep anesthesia (6% desflurane), the mean spike rate was as low as in S1 (burst-suppression period), however a small portion of neurons showed distinctly high spike rate (Fig. 4B-C). Interestingly, the firing pattern of neural population was asynchronous, similar to wakefulness (Fig. 5). Together with the relatively high EMG activity (Fig. 3F-G), this suggested that S3 could be considered a paradoxically aroused state. It can be surmised that there was at least a theoretical possibility of transient awareness in this state. A similar proposal has been put forward for the UP states in slow-wave sleep (Destexhe et al., 2007). Given the unexpected nature of this paradoxical state, replication of this finding together with a more systematic behavioral assessment or level of consciousness will offer a more significant clinical implication and advance the general understanding of the neuronal network dynamics.

### Spiking behavior changes monotonically with brain state

It has been widely reported that most properties of large-scale brain activity (EEG, LFP) exhibit a biphasic pattern with anesthetic depth; i.e., an initial increase (decrease) of EEG/LFP variable is followed by decrease (increase) as anesthesia deepens. (Borgeat et al., 1991; Kuizenga et al., 1998, 2001; Lee et al., 2017). In agreement, in our study, the low-frequency dominance of LFP (L/H ratio) first increased then decreased (ignoring S3) as the anesthetic was stepwise withdrawn. Unexpectedly, the spiking behavior did not follow this biphasic pattern; the changes were always monotonic for all spike properties from burst-suppression (S1) to full wakefulness (S5) (again, ignoring S3). The reason for this discrepancy is not known but it may be due to a limitation of measurements at large-scale level. For example, spike rate decreases monotonically as the anesthesia deepens but this cannot be measured directly by EEG or LFP because these measurements mostly reflect synchronous population activity. In addition, the theory of complex system predicts that for a system consisting of many interacting elements such as the neuronal network, an incremental change local variables can lead to abrupt, qualitative change in macroscopic variables; in this case, leading to biphasic macroscopic behavior.

## Conclusions

We identified five distinct brain states during stepwise changes of the anesthetic state. The identified brain states displayed degeneracy in their relationship with anesthetic concentration suggesting the presence of metastable or multistable dynamics with specific, transient patterns of neuronal spiking. A previously unidentified paradoxically desynchronized state was found during deep anesthesia. The synchronously fragmented spiking in anesthesia appears to be a robust signature of the state of unconsciousness.

## Acknowledgements

Research reported in this publication was supported by the National Institute of General Medical Sciences of the National Institutes of Health under award number R01-GM056398 and the Center for Consciousness Science, Department of Anesthesiology, University of Michigan Medical School, Ann Arbor, Michigan, USA. The content is solely the responsibility of the authors and does not necessarily represent the official views of the National Institutes of Health. The authors thank members of the Center for Consciousness Science, University of Michigan Medical School, for valuable comments and Kathy Zelenock, MS for her assistance in laboratory operations and manuscript editing.

## Notes

Conflicts of Interest: None

### Competing Interest Statement

The authors have declared no competing interest.

